# Highly efficient organ-targeting transport through the ventral midline interstitial channels injection: a new development of interstitium

**DOI:** 10.1101/2025.01.03.631270

**Authors:** Xiaojing Song, Xin Gu, Feng Xiong, Shuyong Jia, Shuyou Wang, Guangjun Wang, Yanpin Wang, Weibo Zhang

## Abstract

How to improve the drugs bioavailability has always been a hot and difficult topic. Conventional drug delivery methods, such as oral administration and the blood circulation system, suffer from issues such as low targeting efficiency, low bioavailability, and high side effects. Recently, interstitium has gradually been recognized as an important new organ. Interstitial injection allows small molecules to move through interstitial channels with interstitial fluid flow and be transported to organs, providing new possibilities for drug delivery. In this study, using fluorescein sodium as a drug model, we systematically investigated the distribution of the fluorescein sodium in organs and body surface of the rats after ventral midline interstitial channels injection. In addition, we observed the microstructure of the ventral midline interstitial channels of abdominal wall and the effects of channels ligation on fluorescein sodium delivery and target organs, to explore the mechanism of interstitial injection. We found that fluorescein sodium can be efficiently transported to the uterus and ovaries along the ventral midline interstitial channels, and the parallel fibers and interconnected interstitial spaces of the channels facilitate long-distance transport of solutes. Additionally, the blockage of the transmission through those channels had results in a significant decrease in targeted transport efficiency over long distances, a decrease in oestrogen levels, organ coefficients, and body weights in female rats. Therefore, the ventral midline interstitial channels with free fluid flow may be a potential communication pathway between body surface and organs. These findings have significant value in the development of drug delivery.

## 1. Introduction

Communication and transport are primarily achieved through the circulatory system in the human body. However, during the early stages of evolution or embryonic development, the specific structures responsible for communication pathways have yet to be fully formed. While the role of the interstitium and interstitial fluid remains unclear, they may potentially contribute to this process. Are there any interstitial channels and the interstitial solute transport that can communicate with different tissues and organs in the body? It is an interesting and meaningful topic.

In the 1960s, the interstitial fluid was believed to be in a unique gel phase and was thought to be almost immobile^1^. According to this perspective, exchange of molecules, ions, and water between blood vessels and the interstitial space primarily occurs through diffusion, which has been the prevailing viewpoint^2^. After extensive calculations and verification, Aukland and Reed proposed a two-phase interstitial fluid model, consisting of a gel phase and a free fluid phase^3,4^. Until 2007, a 10 µm bead was observed using confocal reflectance microscopy within the fibrous composition of an *in vitro* fibrin gel, marking the first direct observation of the free fluid phase^5^.

In 2018, Benias et al. discovered extensive fluid-filled spaces within and between living tissues, validating fluid flow through fibrous in the human interstitium^6^. The interstitial continuity within and between organs was verified by monitoring the movement of nonbiological particles^7^. These interstitial spaces may serve as conduits for particle movement, and interstitial fluid flows could provide the necessary medium. Interstitial fluid may also offer complementary lateral connections that cannot exist within the nervous, vascular, and lymphatic systems. They considered that the body-wide network of fluid-filled interstitial spaces has important implications for molecular signalling, cell trafficking, and the spread of malignant and infectious diseases^7^.

The concept of low hydraulic resistance channels along meridians in the interstitium was proposed by Zhang et al.^8,9^ More recently, long-distance free fluid flow with special tracers through interstitium has been visualized in the *Gephyrocharax melanocheir* fish, rats, miniature pigs, and humans^10–14^. Furthermore, the interstitial space where free interstitial fluid flows is referred to as an interstitial channel, and various types of mass transport occur in this channel^13^. Li et al. also observed long-range flow of interstitial fluid in the arterial and venous adventitia, nerves, and cutaneous tissues by injecting magnetic resonance imaging (MRI) tracers into the human body and amputated limbs^15^, thus inferring the existence of a previously unknown fluid pathway outside the circulatory system. Han^16^ et al. found intrathecal space injection of limbs significantly increased the drug concentration in the lung interstitial fluid than that in oral administration.

Although the essential role of interstitium and interstitial fluid flow in cellular biology, tissue morphogenesis, and tissue engineering is increasingly being recognized^17–20^, it has been paid little attention to their use as a substance transport, especially as a drug delivery pathway. Oral administration often has low bioavailability^21^, intravenous injection can pose the risk of systemic toxic or side effects^22^. Although new materials have been developed, literature research from 2005 to 2015 reported that a median of 0.7% of the injected dose of the nanoparticles reached the tumour^23^. Whereas, in the interstitium, interstitial channels and free interstitial fluid flow will provide a green pathway for long-distance transport of small molecules. It has profound value in improving drug bioavailability, reducing adverse reactions. Interstitial injection, a novel drug delivery mode studied in the past decade, offers new possibilities for drug delivery^24–26^. By injecting drugs into the interstitial space, they can be delivered along low hydraulic resistance channels, achieving targeted delivery to specific organs or tissues. In previous studies, we found an interstitial channel for interstitial fluid flow along the ventral midline in rats using fluorescein sodium tracer^27^. In this study, we aim to further explore the transmission characteristics of small molecules caused by the interstitial fluid flow through the ventral midline interstitial channel. To explore the possibilities of the ventral midline interstitial channel as a new route for drug delivery.

## 2. Materials and methods

### 2.1. Animals and groups

In this study, the female Sprague-Dawley (SD) rats were provided by Beijing Vital River Laboratory Animal Technology Co., Ltd. (Beijing, China). All rats were cared according to the feeding standards for experimental animals. All animal experiments were performed with reference to the protocol for animal experiments as defined by the China Academy of Chinese Medical Sciences, Institute of Acupuncture and Moxibustion (CACMA-IAM), Beijing, China. Ethical approval for this study was granted by the ethics committee of CACMA-IAM.

Forty-eight female rats, aged 4 weeks and weighing 100 g each, were randomly assigned to three groups: the ventral midline interstitial channels ligation group, the control ligation group (ligation adjacent to the ventral midline interstitial channels), and the ventral midline group (without ligation). Each group contained 16 rats. The eight female Sprague-Dawley (SD) rats, aged 4 weeks and weighing 100 g each, were assigned to tail vein group. After one week of acclimatization, fluorescein sodium solution was injected into the ventral midline interstitial channels of the ventral midline group to observe its distribution in various organs, including the heart, lungs, spleen, liver, kidneys, uterus, and ovaries. Serum estrogen levels, organ coefficients, and uterine and ovarian histology were then evaluated after four weeks of feeding under standard conditions. Surgical ligation was performed on the low impedance line in the ligation group and next to the ventral midline of the abdomen in the control ligation group. One week after ligation, fluorescein sodium solution was again injected into the ventral midline interstitial channels to observe its distribution in the organs, and various indicators were measured after four weeks of feeding and access to water under standard conditions.

### 2.2. Detection of the low impedance point of the ventral midline interstitial channels

The rats’ ventral skin was shaved for subsequent experiments. After complete anesthesia with isoflurane (2-3% in oxygen) delivered through an animal anesthesia unit (Matrx Company, Midmark Animal Health, Versailles, OH, USA), they were positioned on the experimental table in a supine position. A meridian locator (WQ6F30, Beijing Donghua Electronic Instrument Factory, Beijing, China) was used to identify the low impedance points on the abdominal skin. A detection electrode was slowly moved along the ventral midline of the abdomen to measure impedance. If the current of the meridian locator significantly increased, the point was marked as a low impedance point. Multiple low impedance points were marked along the ventral midline to form a line.

### 2.3. Ligation of the ventral midline interstitial channels and the tissues adjacent to the ventral midline interstitial channels

Under anesthesia with 3% isoflurane (2-3% oxygen) delivered via an anesthesia unit (Matrx Company, Midmark Animal Health, Versailles, OH, USA), the abdominal skin of the rats was disinfected with 75% alcohol. The low impedance line along the ventral midline between the xiphoid process and external genitalia was divided into five equal parts. The site one-fifth of the distance from the xiphoid process was designated as point A and utilized as the injection site. The midpoint between the xiphoid process and the external genitalia, approximately 0.5 cm below point A, was designated as point B and used as the ligation site in the ligation group. The needle insertion and withdrawal points for ligation were positioned 2.5 mm to the left and right of the center, respectively, using point B as the center. Skin and tissue under the skin at a depth of approximately 2-3 mm were ligated perpendicular to the low impedance line of the ventral midline. Point C, located approximately 1 cm horizontal from point B, was designated as the ligation site in the control ligation group, utilizing the same method as the ligation group (Fig. 1B).

**Figure 1.**
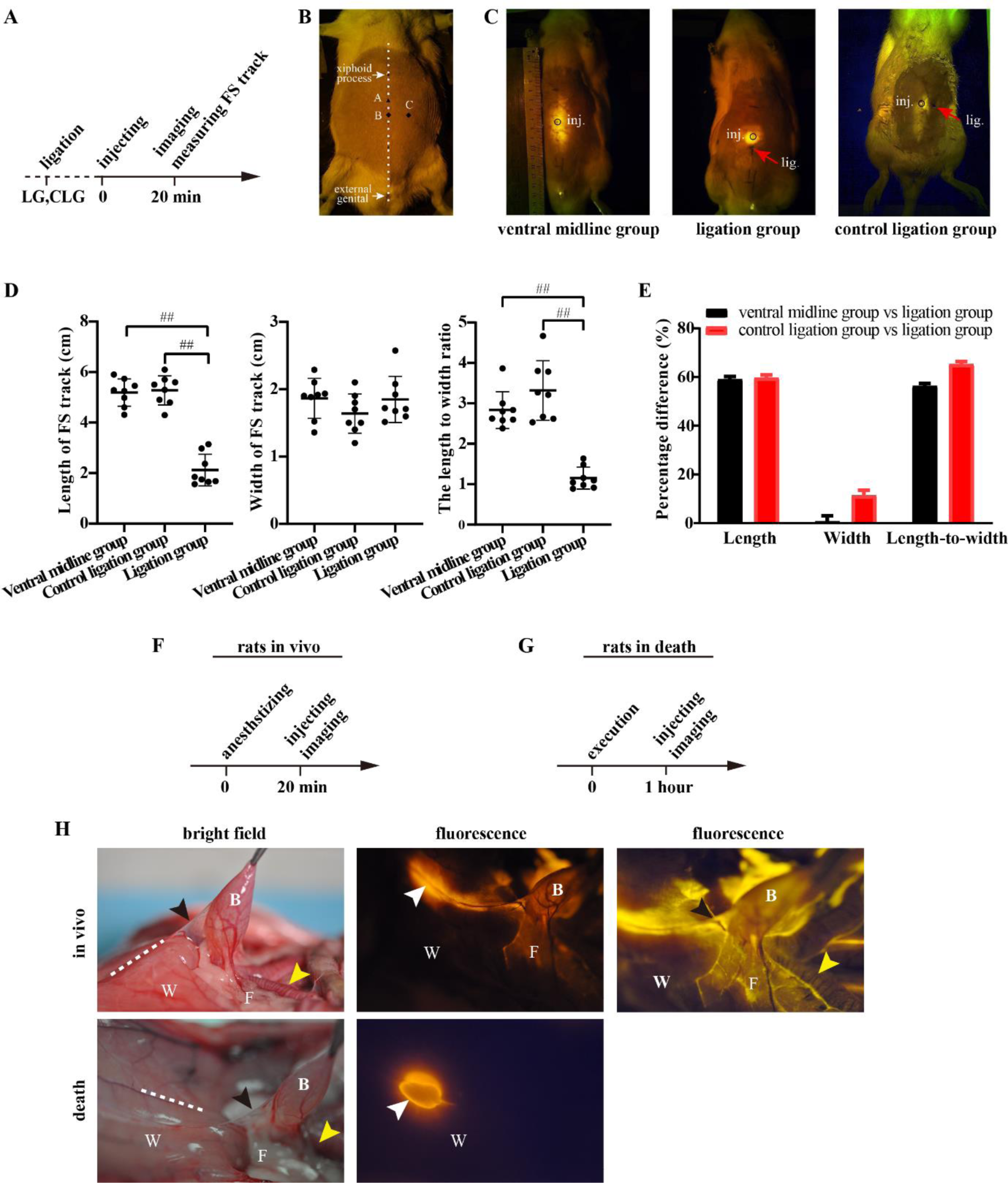
Comparison of fluorescein sodium migration tracks. A) Experiment timeline. The tissue ligation surgery was performed in rats of the ligation group (LG) and control ligation group (CLG). Following that, fluorescein sodium injection and fluorescence imaging were performed. B) Schematic diagram showing the location of the fluorescein sodium injection point (A: on the ventral midline) and ligation point (B: on the ventral midline, C: approximately 1 cm adjacent to the ventral midline) in the rat abdomen. ▴, fluorescein sodium injection point;, ligation point. C) Fluorescein sodium migration was imaged by fluorography at 20 min after fluorescein sodium injection. Fluorescein sodium was transported along the ventral midline of the skin in the ventral midline group without ligation; in the ligation group, fluorescein sodium was concentrated around the injection point and did not cross the ligation point; and in the control ligation group, fluorescein sodium was transported along the ventral midline of the skin. FS: fluorescein sodium; inj., fluorescein sodium injection point; lig., ligation point. D) At 20 min after fluorescein sodium injection into the ventral midline, the length and width of the fluorescein sodium migration track were measured, and each experiment was performed at least two times independently (n = 8). The length-to-width ratio was calculated for each rat. Each dot corresponds to the mean length, width, and length-to- width ratio of the fluorescein sodium migration tracks. The mean and the standard deviation for each group are presented. ^#^ *p* < 0.05 and ^##^ *p* < 0.01, as determined by one-way analysis of variance (ANOVA) with Tukey’s test in comparison to the ligation group. E) The percentage difference in the length, width, and length-to-width ratio between the two groups was calculated as follows: (Mean*group1*/Mean*group2* -1) × 100%. Compared to the ventral midline group and the control ligation group, the ligation group exhibited a shorter migration length of fluorescein sodium and a smaller length-to-width ratio. However, there was minimal disparity in width between the groups. F) Experiment timeline. After sufficient anesthesia, fluorescein sodium injecting and fluorescence imaging were performed simultaneously on the rats *in vivo*. G) Experiment timeline. After 1 hour of the rat death, fluorescein sodium injecting and fluorescence imaging were performed simultaneously. H) At 20 minutes after injecting fluorescein sodium into point A, the fluorescein sodium transported along the ventral midline (dashed line) of interior abdominal wall (W) towards the median umbilical ligament (black arrows), fatty tissue (F) surrounding bladder (B), and periexternal uterus envelop (yellow arrows) of rats *in vivo*. In the death rats, the fluorescein sodium persisted at the injection site (withe arrows), displaying no long-distance migration on the interior abdominal wall after injection (n = 3).

### 2.4. Acquisition and characterization of the fluorescein sodium migration track

Under anaesthesia with 3% isoflurane, all rats were injected with 100 μl of fluorescein sodium (1%) (0.1 ml/kg) (Guangxi Wuzhou Pharmaceutical Co., Ltd., Wuzhou, China) at point A. The rats were then returned to the rearing cage. At 20 min after injection, fluorescein sodium migration images were collected by fluorography (Canon 5D2 camera; optical filter, 570 nm; laser sensor power, 0.5 mW; and excitation wavelength, 455 nm). The length of the fluorescein sodium migration track along the ventral midline and the width perpendicular to the ventral midline were measured, and the length-to-width ratio was calculated. The percentage difference in the length/width/length-to-width ratio between the two groups was calculated as follows: (Mean*group1*/Mean*group2* -1) ×100%. In addition, three rats were performed U-shaped abdominal incisions under deep anaesthesia with 20% urethane (1.35 g/kg; Sigma-Aldrich (Shanghai) Trading Co., Ltd., Shanghai, China). The fluorescein sodium (1%, 25μl) was injected into point A. Then, the fluorescein sodium distributes in the abdominal wall and organs were captured with images by fluorography. The other three rats were killed by an intraperitoneal overdose of 20% urethane. After one hour of death, the rats were operated the aforementioned performances.

### 2.5. Analysis of the fluorescein sodium distribution in organs

Under 3% isoflurane anesthesia, some rats received a 100 μl injection of 1% fluorescein sodium at point A, the other rats received intravenous injection of 100 μl fluorescein sodium through the tail vein. They were sacrificed 20 min later by cervical dislocation. The heart, lungs, liver lobes, spleen, kidneys, uterus, and ovaries were immediately excised and placed in a dark recording chamber of a Carestream In-Vivo Imaging System FX PRO. *In vivo* fluorescence images of the tissues were obtained with an excitation wavelength of 470 nm, an emission wavelength of 535 nm, and a 10-second exposure time. Carestream In-Vivo FX PRO Acquire software was used to analyze the fluorescence intensity of the organs. Additionally, each organ was cut into pieces and soaked in 5 ml of cold ultrapure water to separate the fluorescein sodium from the samples. After 12 hours, the soaking solution was centrifuged at 3166 g and 4°C for 10 minutes to obtain the supernatant, which was then detected and analyzed using a Thermo VARIOSKAN microplate reader and SkanIt 2.4.3 software. The uptake rate of fluorescein sodium in each organ was calculated as follows: uptake rate = [(*C*organ/*C*injection)100%]/*W*organ, *C* is the content of fluorescein sodium, *W* is the weight of organ.

### 2.6. Measurement of the weight and organ coefficients

Each rat was weighed weekly, one week before the model was created and every week from the first to the fifth week after modelling. All rats were deeply anaesthetized with 20% urethane Trading Co., Ltd., Shanghai, China), and samples of blood were taken from the abdominal aorta. The rats were then euthanized by cervical dislocation, and the uterus, ovaries, liver, spleen, and kidneys were immediately removed and washed with normal saline solution. Adhering fat and fascial tissues were carefully removed from the surface of the organs, which were then dried using filter paper. The organs were weighed using an electronic balance. The organ coefficients were calculated as follows: organ coefficient = organ weight/body weight × 1000‰.

### 2.7. Observation of the histology of the ventral midline interstitial channels *in vivo* and *in vitro*

After obtaining images of the fluorescein sodium migration track, under anaesthesia with 3% isoflurane, a transverse incision perpendicular to the ventral midline (approximately 1 cm) was made in the skin of the migration track. The probe of an *in vivo* confocal laser endomicroscopy imaging system (ViewnVivo FIVE2, OptiScan, Austria) was placed vertically on the surface of the ventral midline on the abdomen. The major parameters of the device were as follows: Class 3R semiconductor laser; excitation wavelength, 488 nm; laser power, 10-1000 μW; point scanning mode; controller connection mode, optical fibre coupling; and field of view, 475 μm× 475 μm. The scanning depth (μm) and laser power (μW) were adjusted to achieve a clear structure. Images of subcutaneous tissue along the ventral midline were captured at different depths, and the probe was then moved perpendicularly to the midline at intervals of 0.5 mm to obtain additional images. The microstructure of the tissue in the image field was analyzed. Then each rat was sacrificed by an intraperitoneal overdose of 20% urethane, frozen, and embedded in gelatin. Transverse sections were taken along the fluorescein sodium migration track at intervals of 5 mm (n = 3), and each section was subjected to fluorescence and brightfield imaging using a Canon 5D2 camera with an optical filter at 570 nm, laser sensor power of 0.5 mW, and excitation wavelength of 455 nm. Each transverse section of the migration track was then cryosectioned at 15 μm and sealed after air-drying in the dark. The morphological structure of the tissues was observed using an Olympus BX53 fluorescence microscope with a CCD OLYMPUS DP74 camera and analyzed with CellSens Dimension software.

### 2.8. Histological and Immunohistochemical staining

Continuous slices of the fluorescein sodium migration track (n = 3), the abdominal wall along the ventral midline, ligated tissue (0.5 cm × 1 cm × 0.3 cm) along the ventral midline, uterine tissues, and bilateral ovarian tissues were cut, fixed, and embedded in paraffin to create 5-μm- thick tissue sections. The sections of the uterus and ovaries were stained with HE solution (G4520, Beijing Solarbio Science & Technology Co., Ltd.). Sections of the ventral midline and ligated tissue were stained with Masson solution (G1350, Beijing Solarbio Science & Technology Co., Ltd.). Sections of the subcutaneous fascia on and adjacent to the ventral midline were stained with collagen fiber and elastic fiber staining Kit (EVG-Victoria Method; G1595, Beijing Solarbio Science & Technology Co., Ltd.), an acetylcholinesterase stain solution (copper ferricyanide method; G2110, Beijing Solarbio Science & Technology Co., Ltd.) and toluidine blue O solution (G3661, Beijing Solarbio Science & Technology Co., Ltd.). Some sections were deparaffinized and washed with 0.01 mM phosphate buffered saline (pH 7.4). The antigen retrieval was done by pretreating the sections with citrate buffer of pH 6 for 20 min. Endogenous peroxidase was blocked with 3 % hydrogen peroxide for 8 min at 37 °C. Sections were blocked with 3 % bovine serum albumin for 30 min at room temperature, then incubated overnight at 4 °C with hyaluronic acid binding protein 1 (HABP-1) (1:100, 24474- 1-AP, Proteintech Group, Inc., USA). After being washed with phosphate buffered saline, the sections were incubated with horseradish peroxidase labeled goat anti-rabbit IgG (1:1000, ab205718, Abcam, Cambridge, UK) at 37 °C for 50 min, and 3,3’-diaminobenzidine tetrahydrochloride (DAB, SW1020, Beijing Solarbio Science & Technology Co., Ltd.) was used as a chromogen. The histomorphological changes in these tissues were observed in each group.

### 2.9. Detection of the serum hormone levels

In the fifth week post-ligation, 3-5 ml of blood was collected from rats in the ligation and control ligation groups, as well as the ventral midline group. The blood was centrifuged (4°C, 20,000 g) to obtain the supernatant. The serum E2, FSH, and LH levels were determined using an E2 enzyme-linked immunosorbent assay (ELISA) kit (Beijing Sino-UK Institute of Biological Technology, Beijing, China), LH ELISA kit (Beijing Sino-UK Institute of Biological Technology, Beijing, China), and FSH ELISA kit (Beijing Sino-UK Institute of Biological Technology, Beijing, China), respectively, following the manufacturer’s protocols. A Thermo VARIOSKAN microplate reader and SkanIt 2.4.3 software (Thermo Scientific, Inc., MA, USA) were used for detection and analysis.

### 2.10. Statistical analysis and reproducibility

The data were processed using SPSS 20.0 statistical software. Normally distributed data were subjected to the homogeneity of variance test. Data with equal variance were analysed by one-way analysis of variance (ANOVA), and Tukey’s test was used for pairwise comparisons between groups. All statistical analyses were two-sided; differences were considered significant if ^#^*P* < 0.05 and ^##^*P* < 0.01.

## 3. Results and discussion

### 3.1. Fluorescein sodium migration *in vivo*

Twenty minutes after injecting fluorescein sodium into the low hydraulic resistance channel at the ventral midline, all eight rats in the ventral midline interstitial channels ligation group (ligation group) exhibited circular diffusion spots of fluorescein sodium that were centred around the injection site in their abdomen, with a length-to-width ratio close to 1:1. However, in the ventral midline group, seven out of eight rats exhibited filamentous migration of fluorescein sodium along the ventral midline of the abdomen, with the length of the migrated track ranging from 1.98 to 5.9 cm. Only one rat showed a circular diffusion spot centered on the injection site. For the group with ligation adjacent to the ventral midline (control ligation group), the fluorescein sodium distribution was similar to that of the ventral midline group, and both groups showed filamentous migration along the ventral midline, with a length ranging from 4.3 to 6.5 cm (Fig. 1C). The length and ratio of the migrated track after ligation of the ventral midline were significantly smaller than those in the ventral midline group (*p* = 0.000) and the control ligation group (*p* = 0.000) (Fig. 1D).

Furthermore, the percentage difference in the length of the fluorescein sodium migration track between the ventral midline group and ligation group was 59.07% (95% confidence interval [CI]: 56.68 to 61.47), and that between the ligation group and the control ligation group was 59.73% (95% CI: 57.35 to 62.1). The percentage difference in the length-to-width ratio of the fluorescein sodium migration track between the ligation group and the ventral midline group was 59.42% (95% CI: 57.25 to 61.58), and that between the ligation group and the control ligation group was 65.33% (95% CI: 63.23 to 67.43). In addition, the percentage difference in the width of the fluorescein sodium migration track between the ligation group and the ventral midline group was 0.79 % (95% CI: -5.33 to 3.76); the percentage difference in the width of the fluorescein sodium migration track between the ligation group and the control ligation group was 11.39 % (95% CI: 7.13 to 15.67) (Fig. 1E). Compared to the ventral midline group and the control ligation group, the ligation group showed a significantly shorter migration length of fluorescein sodium and a smaller length-to-width ratio. However, there was little difference in width.

In addition, it was noted that fluorescein sodium was transferred along the ventral midline of interior abdominal wall, passed through the median umbilical ligament to the bladder’s adjacent fatty tissues within 20 minutes post injection. It had also migrated around the uterus, anatomically connected with the bladder. Dead rats’ abdominal walls were injected with fluorescein sodium, yet no migration was observed (Figure 1H).

The fluorescein sodium was injected into the low impedance point of ventral midline channel in female rats, the free interstitial fluid mixed with fluorescein sodium migrated along the channels with relative ease. The results suggested that the directional and long-distance free interstitial fluid flow and transport of solute existed in the interstitium. The interstitum is complex and do need to be recognized furtherly. According to the current knowledge, it is considered that the majority of interstitial fluid in the interstitum is a gel phase composed of glycosaminoglycans and barely flowing. However, in recent years, the existence of free interstitial fluid has been confirmed^5–7,10–15^. The interstitial spaces not only were occupied by glycosaminoglycans, but also by free interstitial fluid mainly made up of water. The free fluid flow follows Darcy’s law and is determined by the pressure gradient and hydraulic resistance of the tissue. Free fluid will easily move towards areas with more empty space and lower hydraulic resistance^28^.

### 3.2. Fluorescein sodium targeted distribution in the uterus and ovaries

In vivo fluorescence imaging revealed that the peak fluorescence signal intensity of every organ was attained 20 minutes after the injection of fluorescein sodium into the ventral midline. Among all the organs observed, the uterus and ovaries displayed the highest fluorescence signal intensity, followed by the kidneys and liver, while the fluorescence signals of the remaining three organs were similar (Figure 2A). Quantification of the fluorescence signal intensity revealed that the fluorescence intensity of the uterus was significantly higher in the ventral midline group and the control ligation group than in the ligation group (p = 0.000, p = 0.000). The fluorescence intensity of the ovaries was significantly higher in the ventral midline group and the control ligation group than in the ligation group (p = 0.000, p = 0.000). Additionally, the fluorescence intensity of the ovaries, uterus and kidney were significantly lower in tail vein injection than in the ventral midline group (p = 0.000, p = 0.000) (Figure 2B). Quantification of the fluorescein sodium uptake rate in the organs of the ventral midline group revealed that the fluorescein sodium uptake rates were highest in the uterus, ovaries, and kidneys, followed by the liver, spleen, lungs, and heart. The rats in the control ligation and ligation groups exhibited similar patterns to those in the ventral midline group. However, in the ligation group, there was an obvious reduction in fluorescein sodium uptake rates in the uterus, ovaries, and kidneys. But the fluorescein sodium uptake rates of the uterus and ovaries were lowest in the tail vein group. It was 5 times higher in the midline group than that in the tail vein group. The fluorescein sodium uptake rates in the uterus (p = 0.006, p = 0.014), ovaries (p = 0.013, p = 0.024) and kidneys (p = 0.001, p = 0.001) were significantly higher in the ventral midline group and the control ligation group than in the ligation group. The fluorescein sodium uptake rates in the uterus (p = 0.000) and ovaries (p = 0.000) were significantly higher in the ventral midline group than in the tail vein group. Furthermore, slight variations were observed in the uptake rates of different organs within the groups (Figure 2C).

**Figure 2.**
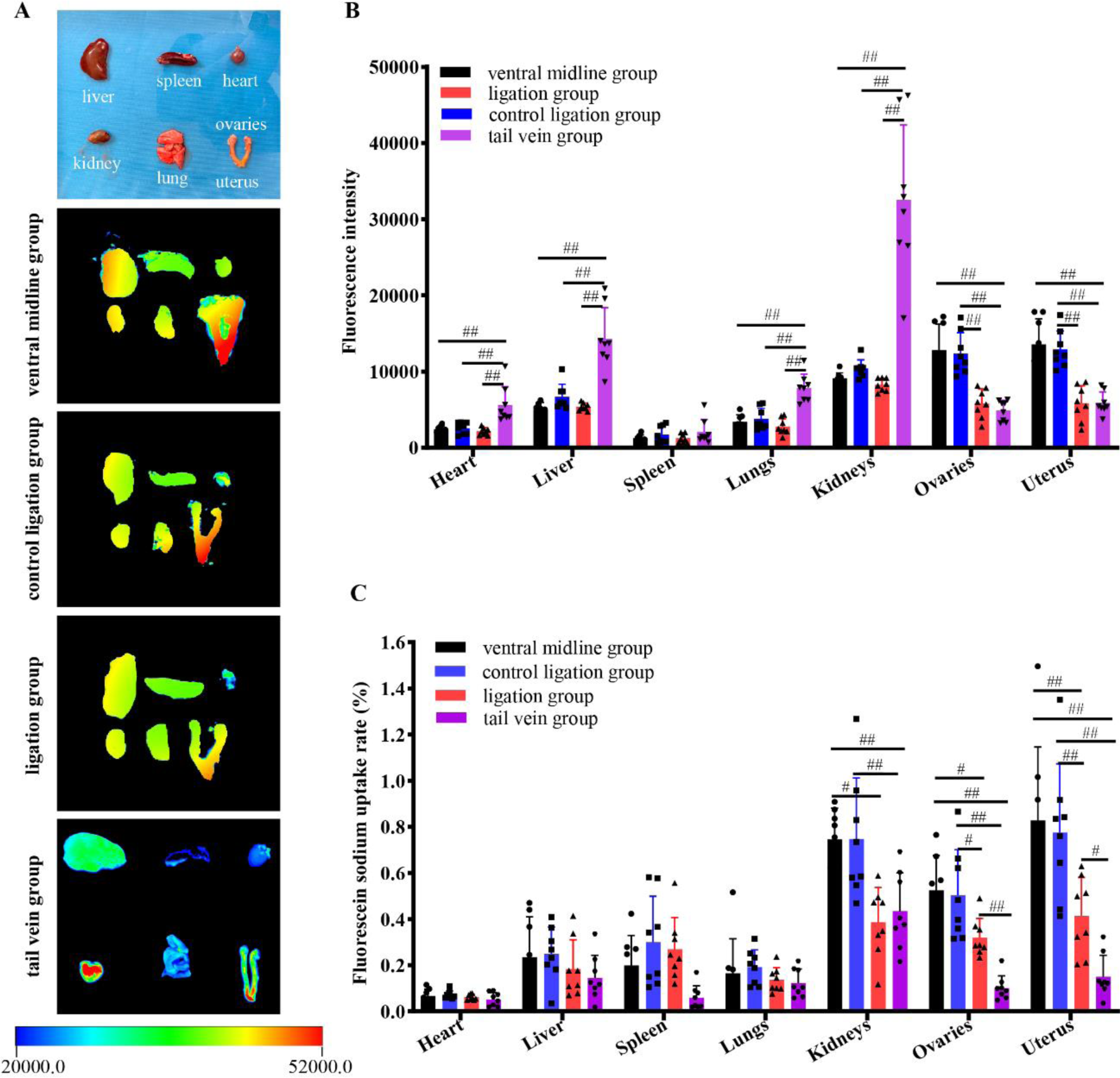
**Organ-based distribution of fluorescein sodium in each group**. A) Fluorescence imaging of the organs was captured at 20 min after fluorescein sodium injection (n = 8). The colour scale (at the bottom) ranges from cool to warm, representing low to high fluorescence signal intensity. In each group, the highest fluorescence intensity was observed in the uterus ovaries and kidneys, and the fluorescence intensity of other organs was similar. Additionally, the fluorescence intensity of the uterus and ovaries was lower in the tail vein group compared to the other groups. B) The fluorescence intensity of the ovaries and uterus was significantly lower in the ligation group than in the ventral midline group and the control ligation group. The fluorescence intensity of the ovaries and uterus was lowest in the tail vein group (n = 8). Each dot corresponds to the mean fluorescence intensity of each organ extracted from fluorescence images; each experiment was performed at least two times independently. The data are presented as the means and standard deviations for each group. C) Comparison of the fluorescein sodium uptake rate among the organs (n = 8). The fluorescein sodium content of each organ was extracted from the optical density of the organ soaking solution; each experiment was performed at least two times independently. The fluorescein sodium uptake rate was calculated for each organ. Each dot corresponds to the mean fluorescein sodium uptake rate. The data are presented as the means and standard deviations for each group. # p < 0.05 and ## p< 0.01, as determined by one-way analysis of variance (ANOVA) with Tukey’s test in comparison to the ligation group.

The above results showed that fluorescein sodium not only was moved along the ventral midline of the skin, but also was transported to the viscera in female rats, with a targeting distribution in the uterus, ovaries, and kidneys. Conventional drug delivery methods, such as oral administration and the blood circulation system, suffer from issues such as low targeting efficiency, low bioavailability, and high side effects. The efficiency of ventral midline interstitial channels injection to transfer to the uterus and ovaries is much higher than that of tail vein injection, which is more than 5 times that of tail vein injection. This is very meaningful for drug targeted delivery.

Meanwhile, the transportation of fluorescein sodium to the uterus and overies was hindered after physically blocking in the ventral midline interstitial channels. However, the obstruction of the ventral midline interstitial channels does not significantly affect the uptake of fluorescein sodium in organs with high blood content, such as the heart, liver, spleen, and lungs, leading us to hypothesize that the free interstitial fluid, mixed with fluorescein sodium in the ventral midline interstitial channels, may be transported through the unique connective tissue networks of the peritoneum (mainly at the bottom of the pelvic cavity) to reach abdominal organs. Studies have shown that the peritoneum, composed of fibrous connective tissue, may serve as a pathway for interstitial fluid transmission and communication between organs and the body^29^. Only the upper part of the bladder is attached to the abdominal wall at the midline of the abdominal wall by ligaments below the diaphragm, while the pubocervical ligament connects the uterus and bladder to the abdominal wall (Figure 1H). Therefore, the interstitial fluid mixed fluorescein sodium in the ventral midline interstitial channels would be preferentially transported to the outer membrane of the uterus, ovaries and bladder, which is also known as the visceral peritoneum through those interconnected ligaments or mesangium. However, the dynamics of free interstitial fluid flow in ventral midline interstitial channels may still be related to the circulatory system, because the long-distance transport did not appear after fluorescein sodium injected into the interstitial channels of dead animals. The high uptake rate of uterus and ovaries resulting from the ventral midline interstitial channels transport is of great significance for the drug delivery.

### 3.3. Weight and organ coefficients change after ligating the ventral midline interstitial channels

In the first and third weeks, the rats in the ligation group demonstrated significantly lower weight compared to those in the ventral midline group during the corresponding period (p = 0.000, p = 0.02). In the third week, the weight of the rats was significantly lower in the ligation group than in the control ligation group (p = 0.01). No significant differences in weight were detected among the groups during the remaining time intervals (Figure 3B).

**Figure 3.**
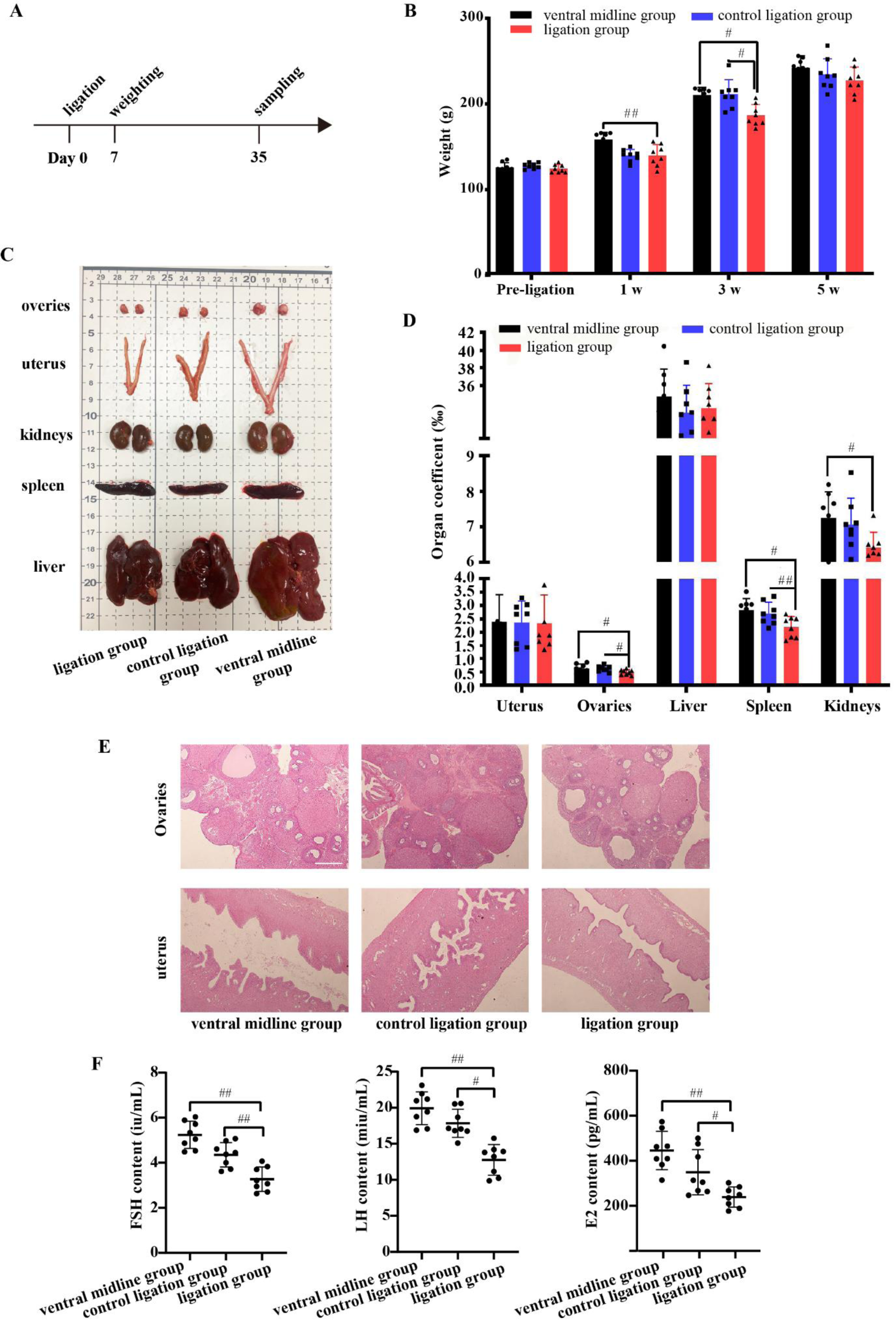
Differences in weight and organ coefficients of rats among the groups. **A)** Experiment timeline. Rats were weighed 1 week before and 4 weeks after ligation surgery. In the fifth week, the rats were euthanized and sampled. B) Comparison of body weight of each group of rats. Following ventral midline ligation, there was a noticeable decrease in the weight of rats, with the ligation group exhibiting significantly lower weight than the ventral midline group after 1 and 3 weeks. Furthermore, at the 3-week mark, the weight of rats in the ligation group was significantly lower than that of rats in the control ligation group (n = 8). Each rat’s weight was measured at least twice independently prior to ventral midline interstitial channels ligation and again at 1, 3, and 5 weeks after ligation. Each dot corresponds to the mean weight of one rat. The data are shown as the means and standard deviations for each group. C) At 5 weeks, rats in the ligation group displayed larger uterus, ovaries, liver, spleen, and kidneys than rats in the control and control ligation groups (n = 8). D) After ventral midline ligation, the organ coefficients of the ovaries, spleen and kidneys significantly decreased (n = 8). The organ weight was measured five weeks after ligation, and each measurement was performed at least two times independently. The organ coefficients were calculated as follows: Organ coefficient = organ weight/body weight × 1000‰. Each dot corresponds to the mean organ coefficient of one rat. The data are shown as the means and standard deviations for each group. # p < 0.05 and ## p < 0.01, as determined by one-way analysis of variance (ANOVA) with Tukey’s test in comparison to the ligation group. E) Haematoxylin and eosin (HE) staining of the ovaries and uterus of rats in each group (×4 magnification, n = 8). The images in the top row showed the ovary, while those in the bottom row showed the uterus. There was no noticeable alteration in the morphology of either the ovaries or uterus following ligation of the ventral midline interstitial channels. Scale bar, 500 μm. F) After ventral midline ligation, the serum oestradiol (E2), luteinizing hormone (LH) and follicle-stimulating hormone (FSH) levels in the rats were significantly decreased (n = 8). The E2, LH, and FSH levels were calculated from the optical density of serum from each rat; each experiment was performed at least two times independently. Each dot corresponds to the mean E2/LH/FSH value of one rat. The data are shown as the means and standard deviations for each group. # p < 0.05 and ## p < 0.01, as determined by one-way analysis of variance (ANOVA) with Tukey’s test in comparison to the ligation group.

A physical map of the organs showed that 5 weeks after ligation, the ovaries, uterus, and kidneys showed a decrease in size compared to the ventral midline group and the control ligation group (Figure 3C). A statistical analysis of the weighs revealed that the organ coefficients of the ovaries were significantly lower in the ligation group than in the ventral midline group (p = 0.032) and the control ligation group (p = 0.042), and the organ coefficient of the kidneys was significantly lower in the ligation group than in the ventral midline group (p = 0.018). In addition, the organ coefficient of the spleen was significantly lower in the ligation group than in the control ligation group (p = 0.024) and the ventral midline group (p = 0.006) (Figure 3D). Aberrant organ development and alterations in body weight were noted following ligation of interstitial channels along the ventral midline in female rats.

### 3.4. Morphology characteristic of the uterus and ovaries after ligating the ventral midline interstitial channels

In the ventral midline group, the ovaries were oval, reddish, shiny, and full and exhibited multiple protruding nodules of different sizes on their surface. HE staining of sections revealed that the cortical layer appeared thick and consisted of numerous follicles showing varied stages of development. The loose connective tissue structure was uniform in the medulla, which contained abundant blood vessels. No noteworthy variations in ovarian gross morphology were observed between the ligation group and the control ligation group or the ventral midline group. There was no significant change in the count of follicular cells at all cortical layer levels, with the interstitium showing abundant blood vessels (Figure 3E). In the ventral midline group, the uterus was pink, the surface was smooth, there were no signs of hydrops or exudation in the uterine cavity, and there were no observed adhesions to the surrounding tissues. HE staining revealed that the mucosa, myometrium, and serous membrane of the uterine cavity were normal and without any anomalies. There were no discernible differences in the uterine morphology between the ligation group, control ligation group, and ventral midline group. No adhesions were observed in the lumen (Figure 3E).

### 3.5. Serum oestrogen levels after ligating the ventral midline interstitial channels

The sexual maturity period of female Sprague‒Dawley (SD) rats begins approximately 60 days after birth. At the 5th weeks after ventral midline interstitial channels ligation in female SD rats, the serum oestradiol (E2) (p = 0.014 vs ventral midline group), luteinizing hormone (LH) (p = 0.000 vs ventral midline group), and follicle-stimulating hormone (FSH) levels (p = 0.000 vs ventral midline group) were significantly lower in the ligation group than in the ventral midline group, and the FSH level was significantly lower in the control ligation group than in the ventral midline group (p = 0.000). The serum E2 (p = 0.021 vs control ligation group), LH (p = 0.000 vs control ligation group), and FSH levels (p = 0.004 vs control ligation group) were significantly decreased in the ligation group (Figure 3F). After ventral midline interstitial channels ligation, the serum E2, LH, and FSH levels and the organ coefficient of the ovaries were significantly decreased in female rats.

Recently, interstitium has gradually been recognized as an important new organ. The blockage in the ventral midline interstitial channels not only hindered the transportation of fluorescein sodium to the uterus and overies through interstitial fluid flow, but also ultimately had an impact on reproductive function (serum E2, FSH, and LH levels) and overall development (weight) in female rats. This finding suggests that the ventral midline interstitial channels and the solute transmission in the channels would be involved in physiological activities of the uterus and ovaries, the ventral midline interstitial channels would be developed as a special communication and transport pathway targeted to the uterus and ovaries independent of the circulatory system.

### 3.6. Histological characteristics of migration track in the ventral midline of the abdomen

In the *in vivo* confocal laser endomicroscopy images, the illuminated areas represent connective tissues or interstitial spaces filled with interstitial fluid marked by fluorescein sodium, while dark areas correspond to other tissues such as blood vessels, cells and muscle fibres. Through confocal laser endomicroscopy *in vivo*, we observed the subcutaneous tissue on the fluorescein sodium migration track. We observed a large amount of connective tissue, small blood vessels, and lymphatic capillaries present in the subcutaneous fascia of the ventral midline, with most fibres arranged in parallel. The interstitial spaces around the fibres were filled with a significant amount of interstitial fluid. At approximately 0.5 cm from the ventral midline, collagen fibres were arranged in a reticular pattern, many dense muscle fibres were present at a deeper level, and only thin white lines distributed in the muscle fibres. The brightness of those areas was less than that of the ventral midline under the same scanning parameters. We concluded that a larger volume of free interstitial fluid was present along the ventral midline than that of outside (Figure 4A).

**Figure 4.**
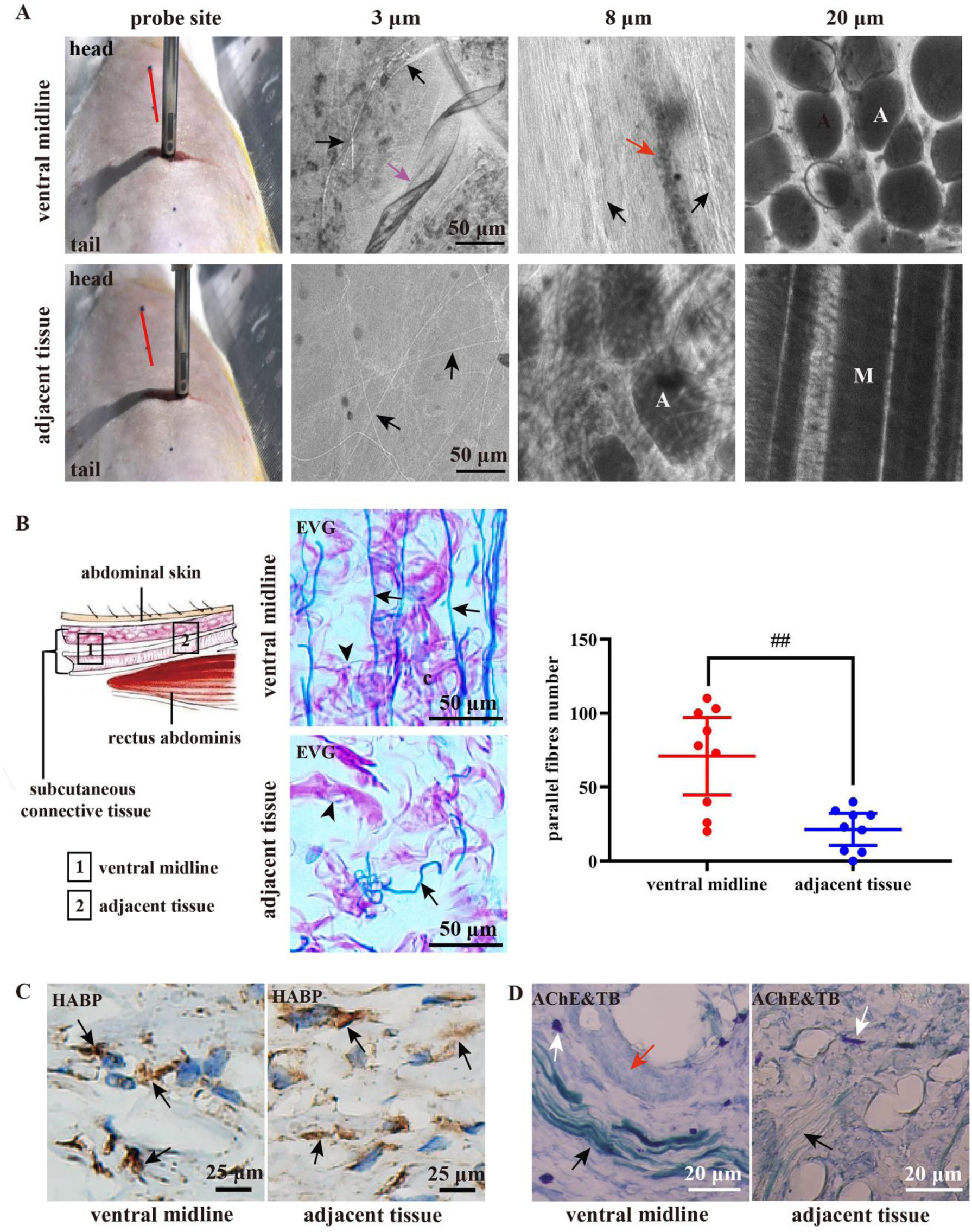
Histology of the subcutaneous fascia on the fluorescein sodium migration track along the ventral midline *in vivo* and *in vitro*. A) The confocal laser endomicroscopy probe was placed on the subcutaneous fascia of the ventral midline (red line) and approximately 0.5 cm away. The histological characteristics at a depth of 3 μm, 8 μm, 20 μm observed by the probe were displayed. The mount of fluorescein sodium indicates the brightness. Free fluid (marked by fluorescein sodium) filled the loose connective tissue from shallow to deep layers. Many parallel fibres (black arrows), blood vessels (red arrow) and lymphatic capillaries (purple arrow) were observed in the subcutaneous fascia of the fluorescein sodium migration track along the ventral midline (upper panel). In the same depth 0.5 cm adjacent to ventral midline, the fibres were arranged in a reticular pattern (black arrows), and adipocytes (A) and muscle fibres (M) were predominantly distributed (×1000 magnification, n = 3) (lower panel). Scale bar, 50 μm. B) The diagram (left) indicates the location (rectangular marks) of the tissue observed. The images (right) are the subcutaneous fascia stained with Victoria blue and improved Van-Gieson staining solution (EVG, ×40 magnification, n = 3). In the subcutaneous fascia along the ventral midline, many elastic fibres (blue, long arrow) arranged in a reticular pattern and collagen fibres (red, short arrow) interwoven into a network were observed. Scale bar, 50 μm. There were significantly more parallel fibres in the subcutaneous fascia of ventral midline than in the adjacent tissue (The whole field were counted for each of the three tissue layers per specimen, each specimen of one rat, with a total of 9 fields represented per group, n = 3). Each dot corresponds to the number of parallel fibres in one slice. The mean and the standard deviation for each group are presented. ## p < 0.01, determined by one-way analysis of variance (ANOVA) with Tukey’s test in comparison to the adjacent tissue. C) Immunohistochemical staining of hyaluronic acid binding protein 1 (HABP-1) in the subcutaneous fascia (HABP, ×40 magnification, n = 3). HABP-1 staining detected hyaluronic acid localization (brown, arrow) to the surrounding collagenous fibres. Scale bar, 25 μm. (D) Acetylcholinesterase (AChE) and toluidine blue (TB) staining of subcutaneous fascia (AChE & TB, ×40 magnification, n = 3). More cholinergic nerves (black arrow), mast cells (white arrow) and blood vessels (red arrow) were observed in the subcutaneous fascia of the ventral midline than in the tissue adjacent to the ventral midline. Scale bar, 20 µm.

In the subcutaneous fascia of the ventral midline *in vitro*, a significant number of long and thin elastic fibres were arranged in parallel, and the collagen fibres were interwoven into a network, while in the adjacent tissues, elastic fibres are mainly disordered. There were significantly more parallel fibres in the subcutaneous fascia of the ventral midline than that in adjacent tissues (*p* = 0.001) (Figure 4B). The hyaluronic acid was adherent around the skeleton of collagenous fibres in the subcutaneous connective tissue, both in the ventral midline and adjacent area, but it was not much (Figure 4C). Hyaluronic acid is mainly adhered to the skeleton of collagenous fibres rather than full-filled in the interstitial space, which is conducive to the flow of free interstitial fluid. Additionally, it was observed that there were some cholinergic nerves, mast cells, and small blood vessels in the connective tissues. (Figure 4D).

Serial transverse sections of the fluorescein sodium migration track along the ventral midline revealed that it migrated not only along the skin surface but also down to the peritoneum, including the parietal and visceral peritoneum (Figure 5B, C). Fluorescence images showed an extensive interstitial network of subcutaneous connective tissue on the ventral midline between the rectus abdominis on either side. Some intersecting gaps were noted in this interstitial network, and a significant amount of interstitial fluid mixed with fluorescein sodium filled these spaces *in vivo*. These interstitial networks of connective tissue extended continuously along the ventral midline (Figure 5D). Masson staining demonstrated that the subcutaneous connective tissues stained blue on the ventral midline were longitudinally distributed to the parietal peritoneum, and large gaps were observed between the rectus abdominis and the longitudinal connective tissue (Figure 5E). Overall, the parallel fibres along the ventral midline, the vast amount of continuous, bound connective tissue, and interstitial spaces would connect to form interstitial channels, which would be conducive to the flow of interstitial fluid. At the site of ventral midline ligation, the skin, connective tissue on the ventral midline, and surrounding muscles were compressed and adhered, various tissues were clustered together, and no large interstitial gaps were observed (Figure 5F, G).

**Figure 5.**
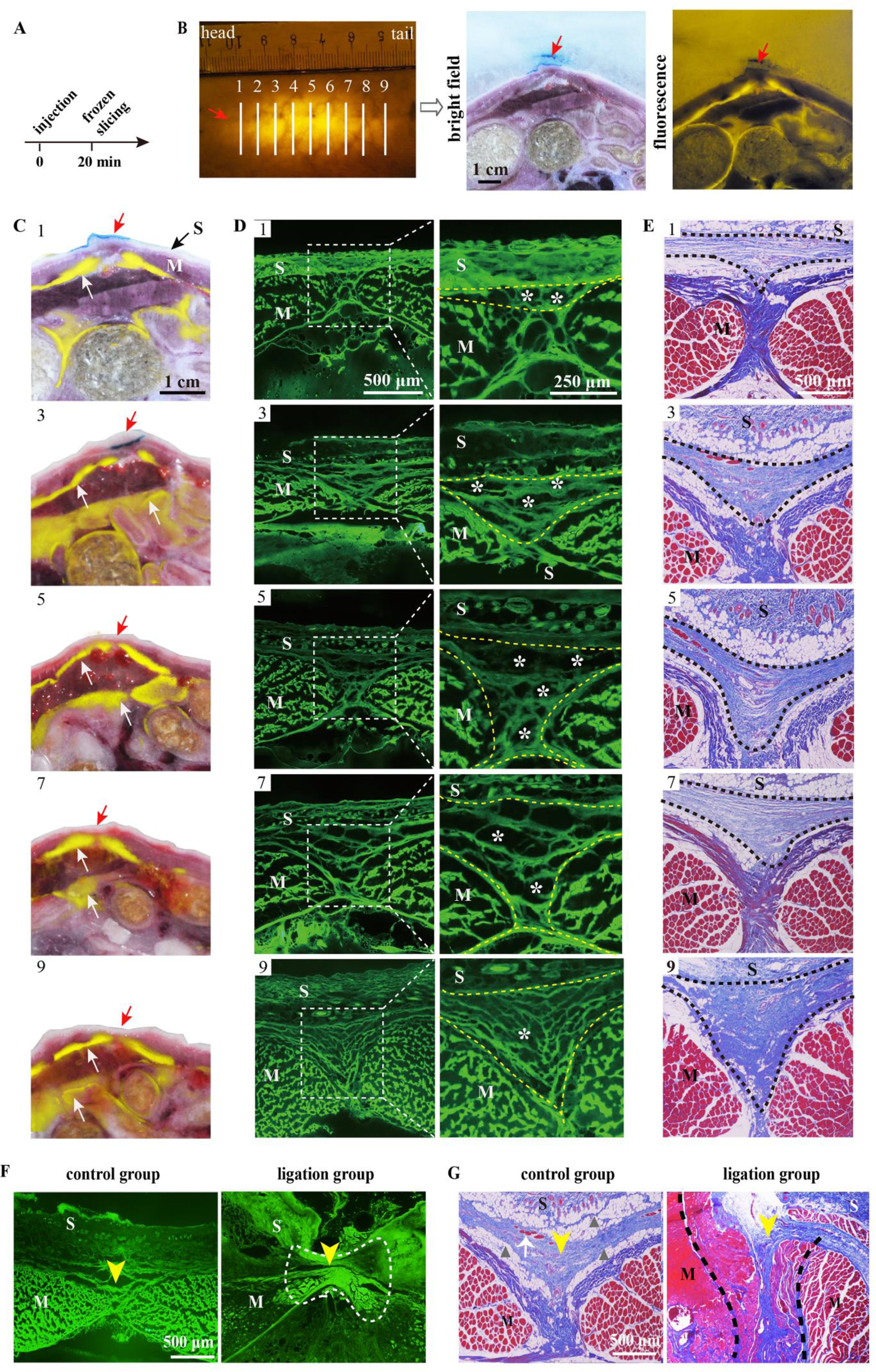
Histology of transverse sections along migration track. A) Experiment timeline. At 20 minutes after fluorescein sodium injection, the animals were euthanized, frozen, and sliced. B) The fluorescein sodium migration track running along the ventral midline (red arrow) was divided into nine parts at 5 mm intervals (n = 3). Transverse sections were then obtained, and both bright field and fluorescence images were recorded. Scale bar, 1 cm. C) Fluorescence signals (white arrows) (Supplementary Figure 1) of each fluorescence image (Supplementary Figure 2) extracted by Photoshop were merged with the corresponding bright field image (Supplementary Figure 3). In this way, the merged images of slices 1, 3, 5, 7, and 9 (B) were obtained, in that order, from top to bottom. The merged images depict fluorescein sodium migrating across the skin (S) of the ventral midline (red arrow) and being transported to the parietal peritoneum (white arrows) as well as the surface membrane of organs. M, muscle; L, liver; SP, spleen; I, intestine; Scale bar, 1 cm. D) Each of the slices was then cryosectioned at a thickness of 15 μm. The left panels show the morphology of the fluorescein sodium migration track on the fresh ventral midline tissue. In the right panels, a considerable amount of loose connective tissue (dashed line) at the subcutaneous level on the ventral midline between the rectus abdominis muscles (M). Many interstitial spaces (* marks) filled with interstitial fluid in the connective tissue were observed in vivo. These spaces were dark due to fluorescent fluid having drained during processing, but the tissue structures were still stained with residual fluorescein. And between the connective tissue and the rectus abdominis, a distinct boundary was observed (×4 magnification, n = 3). Scale bar of left image, 500 μm; scale bar of right image, 250 μm. E) Masson staining of serial sections revealed the same structure observed in Fig. 3c. The loose connective tissue (dashed line) formed a boundary with the rectus abdominis muscles (M), which were stained purple and distributed from the subcutaneous level to the parietal peritoneum on the ventral midline (×4 magnification, n = 3). Scale bar, 500 μm. F) The fresh-frozen sections showed that the skin (S), subcutaneous connective tissue and rectus abdominis (M) were clustered together in the ligated tissue (dashed line) at the ventral midline (yellow arrow), with no significant interstitial space detected. (×4 magnification, n = 3). Scale bar, 500 μm. G) Masson staining showed the absence of interstitial spaces in the connective tissue on the ventral midline (yellow arrow) and the absence of large gaps (▴ marks) between the connective tissue and rectus abdominis (M). This observation was within the range covered by the ligation line (dashed line) (×4 magnification, n = 3). White arrow, blood vessels. Scale bar, 500 μm.

Levick observed the existence of collagen fibre peripheral spaces ranging from 1.4 to 1.7 nm, allowing free fluid to flow^2^. And these spaces are collectively referred to as the collagen fibre peripheral space. The parallel alignment of fibres facilitated the formation of interstitial fluid flow pathways^30,31^. Han et al. suggested the fast molecular transport within micro/nanoscale multiphase porous systems was a green pathway different from simple diffusion in soft matter^32^. Histological results could explain why solute transport occurs in ventral midline interstitial channels, which provided structure conditions for interstitial transport. A large amount of free interstitial fluid marked by the water indicator fluorescein sodium full filled in those interstitial spaces. And the parallel fibres and relatively large and continuous spaces along the ventral midline interstitial channels will lead to a low hydraulic resistance. Therefore, the microstructure of the subcutaneous ventral midline enables the creation of multiple interconnected interstitial channels for free interstitial fluid flow.

Interstitial transport is a key area of study in developmental biology^33^. The movement of interstitial fluid within the interstitium is critical for solute transport, intercellular communication^34,35^. This flow can also stimulate blood and lymphatic capillary morphogenesis *in vitro* and lymphatic regeneration *in vivo*^36^. Additionally, it has been found to sustain the functional activity of chondrocytes and osteocytes ^37^, and induce fibroblast differentiation^38^. If free fluid flow is obstructed, it can impact mass transportation and bioinformatic signal transmission, which in turn can affect the homeostasis of the environment, metabolic functions, and organ development along the flow pathway^39–42^. During development, there may be an interaction between free interstitial fluid flow and parallel fibres, resulting in the creation of low hydraulic resistance channels. These channels serve as natural pathways for free fluid flow and play an important role in replenishing vascular circulation, transporting small-molecule signals, and supporting tissue repair.

While the ventral midline was ligated, some small blood and lymphatic capillaries were occluded in the subcutaneous midline, it would reduce the sources of substances and free fluid and disrupt the dynamics of free fluid flow in the interstitial channels. It would lead to weaker solute transportation and bioinformation transmission between these channels and the viscera, especially the uterus and ovaries. The appearance of reduction in the ovarian coefficient and a decrease in estrogen levels of female rats suggested that there was communication between the ventral midline interstitial channels and the uterus, ovaries. The interstitial transport of ventral midline interstitial channel and its potential communication within uterus and ovaries can be used to explore an innovative approach for reproductive disease treatment in the future. Although our data indicated that interstitial fluid mixed fluorescein sodium targeted transport to the uterus and ovaries through interstitial channels. We did not directly observe the dynamic motion of free interstitial fluid mixed with tracer at different depths of the ventral midline interstitial channel in the abdominal wall in real time. In addition, the kinetics and targeting characteristics of drugs or other particles injection through the ventral midline interstitial channels were unknown. These issues necessitate further investigation.

## 4. Conclusion

In conclusion, in the interstitium, the interstitial channels with the free interstitial fluid flow is critical for solute transport and intercellular communication. Our findings add to the emerging literature that soluble substances in free interstitial fluid can have a targeted delivery to the uterus and ovaries through ventral midline interstitial channels, the parallel fibres and the large and continuous fibroskeletal spaces in the ventral midline interstitial channels facilitate the free interstitial fluid flow. These findings are valuable for the development of drug delivery approach for the abdominal viscera and tissues, especially the uterus and ovaries.

## Ethical approval

All animal experiments were conducted on the basis of the Guidelines for Ethical and Regulatory for China Academy of Chinese Medical Sciences, Institute of Acupuncture and Moxibustion (CACMS-IAM), China. The animal ethics committee of CACMS-IAM granted ethical approval for this study. The approval number of the ethics committee was D-2023-01- 10-01. The rats were raised at 25 °C and a relative humidity of 70 ± 5% under 12 h light–dark cycle environment for 1 week before the experiments. In this study, all invasive operations to the experimental rat were done under complete anesthesia with isoflurane (2-3% in oxygen). In addition, to minimize the suffering of rat as much as possible, a clean and comfortable living environment was provided, ensuring sufficient food and water sources. At the end of the experiment, a part of rats was euthanized by an intraperitoneal overdose of 20% urethane, other rats were then euthanized by cervical dislocation. The authors have adhered to the ARRIVE guidelines.

## Author Contributions

W.B. Zhang contributed to the study design, interpreted the data, and co-wrote the manuscript.

X.J. Song wrote the manuscript, interpreted the data, and performed all tissue staining and observations of the ventral midline interstitial channels *in vivo* by confocal laser imaging. X. Gu carried out ligation of the ventral midline interstitial channels, measured the organ coefficients, weights and serum hormone levels, and co-wrote the “Methods” section. F. Xiong contributed to the analysis of the fluorescein sodium migration tracks and production of figures.

S.Y. Jia and S.Y. Wang contributed to performing the animal experiment and acquisition original data. G.J. Wang contributed to interpreted the data and providing references. Y.P. Wang contributed to the study design and editing the manuscript.

## Supporting Information

Supporting Information is available from the authors.

## Acknowledgements

The authors wish to thank Yuqiang Jiang and Rongcheng Han for their excellent guidance on the design of fluorescein sodium tracing experiments. The authors acknowledge support from Yinan Shi for analysis on uptake rate of fluorescein sodium in each organ. Beijing Shengou Technology Co., Ltd. and technical staff are gratefully acknowledged for excellent technical expertise and assistance concerning for histological staining of interstitial tissue. The authors acknowledge for English editing from American Journal Experts (AJE).

## Data Availability Statement

The data generated or analysed during this study are provided in the main paper (Figs. 1-6) and supplementary figures (Supplementary Figs. 1-3). The original data for the graphs can be found in the Supplementary files. All morphological images generated and/or utilized in this study are stored in the workstations of the Department of Scientific Research, Institute of Acupuncture & Moxibustion, China Academy of Chinese Medical Sciences. The data supporting the findings of this study are available from the corresponding author upon reasonable request.

## Additional information Funding

This work was supported by a grant from the National Natural Science Foundation of China (82050006) and the CACMS Innovation Fund (CI2021A03406).

## References

1. Guyton AC, Scheel K, Murphree D. (1966) Interstitial fluid pressure. III. Its effect on resistance to tissue fluid mobility. Circ Res 19(2):412–9. doi: 10.1161/01.

2. Guyton AC, Taylor AE, Granger HJ. (1975) Circulatory Physiology. Dynamics and control of the body fluids. Philadelphia. 1975. (PA:Saunders).

3. Aukland K, Reed PK. (1993) Interstitial-lymphatic mechanisms in the control of extracellular fluid volume. Physiol Rev 73(1):1–78. doi: 10.1152/physrev.1993.73.1.1.

4. Levick JR. (1987) Flow through interstitium and other fibrous matrices. Q J Exp Physiol. 72(4):409–37. doi: 10.1113/expphysiol.1987.

5. Swartz MA, Fleury ME. (2007) Interstitial flow and is effects in soft tissues. Annu. Rev. Biomed. Eng 9:229–256. doi: 10.1146/annurev.bioeng.9.060906.151850. .

6. Benias PC, Wells RG, Sackey-Aboagye B, et al. (2018) Structure and distribution of an unrecognized interstitium in human tissues. Sci Rep 8(1):4947. doi: 10.1038/s41598-018-23062-6.

7. Cenaj O, Allison Douglas HR, Imam R, et al. (2021) Evidence for continuity of interstitial spaces across tissue and organ boundaries in humans. Commun Biol 4(1):436. doi: 10.1038/s42003-021-01962-0.

8. Zhang WB, Tian YY, Li H, et al. (2008) A Discovery of Low hydraulic resistance channel along meridians. J Acupunct Meridian Stud 1(1):20–8. doi: 10.1016/S2005-2901(09)60003-0.

9. Zhang WB, Wang GJ, Fuxe K. (2015) Classic and modern meridian studies, A review of low hydraulic resistance channels along meridians and their relevance for therapeutic effects in traditional Chinese medicine. Evid Based Complement Alternat Med 2015:410979. doi: 10.1155/2015/410979.

10. Wang Z, Zhang WB, Jia SY, et al. (2015) Finding blue tracks in GM fish similar to the locations of acupuncture meridians after injecting Alcian blue. J Acupunct Meridian Stud 8(6):307–13. doi: 10.1016/j.jams.2015.08.007.

11. Song XJ, Zhang WB, Jia SY, et al. (2021) A discovery of low hydraulic resistance channels along meridians in rats. World. J. Acup. Moxib 31(1):22–29.doi: 10.1016/j.imr.2020.100560

12. Gu X, Wang YP, Wang GJ, et al. (2020) In vivo display of low hydraulic resistance channels along conception vessel in rats by photofluorography. Zhen. Ci. Yan. Jiu. 45(3):227–232. doi: 10.13702/j.1000-0607.190692.

13. Xiong F, Song XJ, Jia SY, et al. (2020) Preliminary observation of the migration of sodium fluorescein along meridians in the limbs of mini-pigs. Scientia. Sinica. Vitae. 50(12):1453–1463.

14. Li, T. J. Tang BQ, Zhang WB, et al. (2021) In vivo visualization of the pericardium meridian with fluorescent dyes. Evid. Based. Complement. Alternat. Med 2021:5581227. doi: 10.1155/2021/5581227.

15. Li, H. Y. Wang F, Chen M, et al. (2021) An acupoint-originated human interstitial fluid circulatory network. Chine. Med 134(19):2365–2369. doi: 10.1097/CM9.0000000000001796.

16. Shi YH, Jiao CQ, Lu X, et al. (2022) Rapamycin nanoparticles improves drug bioavailability in PLAM treatment by interstitial injection. Orphanet J Rare Dis 17(1):349. doi: 10.1186/s13023-022-02511-6.

17. Fogh-Andersen N, Altura BM, Altura BT, et al. (1995) Composition of interstitial fluid. Clin. Chem. 41(10):1522–1525.

18. Losada-Barragán M, Umaña-Pérez A, Rodriguez-Vega A, et al. (2019) Proteomic profiling of splenic interstitial fluid of malnourished mice infected with Leishmania infantum reveals defects on cell proliferation and pro-inflammatory response. J. Proteomics. 208:103492. doi: 10.1016/j.jprot.2019.103492.

19. Nin F, Yoshida T, Sawamura S, et al. (2016) The unique electrical properties in an extracellular fluid of the mammalian cochlea; their functional roles, homeostatic processes, and pathological significance. Pflugers. Arch 468(10):1637–49. doi: 10.1007/s00424-016-1871-0.

20. Agnati LF, Marcoli M, Leo G, et al. (2017) Homeostasis and the concept of ’interstitial fluids hierarchy’: Relevance of cerebrospinal fluid sodium concentrations and brain temperature control (Review). Int. J. Mol. Med 39(3):487–497. doi: 10.3892/ijmm.2017.2874.

21. Naeem A, Ming Y, Pengyi H, et al. (2022) The fate of flavonoids after oral administration: a comprehensive overview of its bioavailability. Crit Rev Food Sci Nutr 62(22):6169–6186. doi: 10.1080/10408398.2021.1898333.

22. Lawhn-Heath C, Fidelman N, Chee B, et al. (2021) Intraarterial Peptide Receptor Radionuclide Therapy Using 90Y-DOTATOC for Hepatic Metastases of Neuroendocrine Tumors. J Nucl Med 62(2):221–227. doi: 10.2967/jnumed.119.241273.

23. Wilhelm S, Tavares AJ, Dai Q, et al. (2016) Analysis of nanoparticle delivery to tumours. Nature Reviews Materials 1:1–12. doi:10.1038/natrevmats.2016.14.

24. Thomas SN, Schudel A. (2015) Overcoming transport barriers for interstitial-, lymphatic-, and lymph node-targeted drug delivery. Curr Opin Chem Eng 7:65–74. doi: 10.1016/j.coche.2014.11.003.

25. Shi YH, Jiao CQ, Lu X, et al. (2022) Rapamycin nanoparticles improves drug bioavailability in PLAM treatment by interstitial injection. Orphanet J Rare Dis Sep 17(1):349. doi: 10.1186/s13023-022-02511-6.

26. Liu WT, Cao YP, Zhou XH, et al. (2022) Interstitial Fluid Behavior and Diseases. Adv Sci (Weinh) 9(6):e2100617. doi: 10.1002/advs.202100617.

27. Song XJ, Xiong F, Jia SY, et al. (2021) Observation of Microstructure of Midline Interstitial Channels of the Inner Abdominal Wall in Rat Applying in vivo Confocal Laser Imaging. Acta. Laser. Biology. Sinica 30(5):469–474.

28. Li L, Iskander M. (2022) Visualization of Interstitial Pore Fluid Flow. J Imaging 8(2):32. doi: 10.3390/jimaging8020032.

29. Morellea J, Devuyst O. (2015) Water and solute transport across the peritoneal membrane. Curr. Opin. Nephrol. Hypertens 24(5):434–43. doi: 10.1097/MNH.0000000000000151.

30. Letechipia JE, Alessi A, Rodriguez G, et al. (2010) Would increased interstitial fluid flow through in situ mechanical stimulation enhance bone remodeling? Med Hypotheses 75(2):196–8. doi: 10.1016/j.mehy.2010.02.021.

31. Keller SB, Averkiou MA. (2022) The Role of Ultrasound in Modulating Interstitial Fluid Pressure in Solid Tumors for Improved Drug Delivery. Bioconjug Chem. 2022,33(6):1049- 1056.

32. Feng J, Wang F, Han XX, et al. (2014) A “green pathway” different from simple diffusion in soft matter: fast molecular transport within micro/nanoscale multiphase porous systems. Nano Res 7(3):434–442. doi: 10.1007/s12274-014-0409-z.

33. Zagadou BF, Barbone PE, Mountain DC. (2020) Significance of the microfluidic flow inside the organ of corti. J. Biomech. Eng 142(8):081009. doi: 10.1115/1.4046637.

34. Wiedenmann CJ, Gottwald C, Zeqiri K,et al. (2023) Slow Interstitial Fluid Flow Activates TGF-β Signaling and Drives Fibrotic Responses in Human Tenon Fibroblasts. Cells 12(17):2205. doi: 10.3390/cells12172205.

35. Losada-Barragán M, Umaña-Pérez A, Rodriguez-Vega A, et al. (2019) Proteomic profiling of splenic interstitial fluid of malnourished mice infected with Leishmania infantum reveals defects on cell proliferation and pro-inflammatory response. J. Proteomics 208:103492. doi: 10.1016/j.jprot.2019.103492.

36. Nakada T, Kwee IL. (2019) Fluid dynamics inside the brain barrier: current concept of interstitial flow, glymphatic flow, and cerebrospinal fluid circulation in the brain. Neuroscientist 25(2):155–166. doi: 10.1177/1073858418775027.

37. Cardoso L, Fritton SP, Gailani G, et al. (2013) Advances in assessment of bone porosity, permeability and interstitial fluid flow. J Biomech 46(2):253–65. doi: 10.1016/j.jbiomech.2012.10.025.

38. Nithiananthan S, Crawford A, Knock JC, et al. (2017) Physiological Fluid Flow Moderates Fibroblast Responses to TGF-β1. J Cell Biochem 118(4):878–890. doi: 10.1002/jcb.25767.

39. Jasuja H, Jaswandkar SV, Katti DR, et al. (2023) Interstitial fluid flow contributes to prostate cancer invasion and migration to bone; study conducted using a novel horizontal flow bioreactor. Biofabrication 15(2):025017. doi: 10.1088/1758-5090/acc09a.

40. Xiang J, Hua Y, Xi G, et al. (2023) Mechanisms of cerebrospinal fluid and brain interstitial fluid production. Neurobiol Dis 183:106159. doi: 10.1016/j.nbd.2023.106159.

41. Hirose M, Asano M, Watanabe-Matsumoto S, et al. (2021) Stagnation of glymphatic interstitial fluid flow and delay in waste clearance in the SOD1-G93A mouse model of ALS. Neurosci Res 171:74–82. doi: 10.1016/j.neures.2020.10.006.

42. Gatti V, Gelbs MJ, Guerra RB, et al. (2021) Interstitial fluid velocity is decreased around cortical bone vascular pores and depends on osteocyte position in a rat model of disuse osteoporosis. Biomech Model Mechanobiol 20(3):1135–1146. doi: 10.1007/s10237-021-01438-4.

