## Supplementary Figure 1,Supplementary Figure 2,Supplementary Figure 3 for "Highly efficient organ-targeting transport through the ventral midline interstitial channels injection: a new development of interstitium"

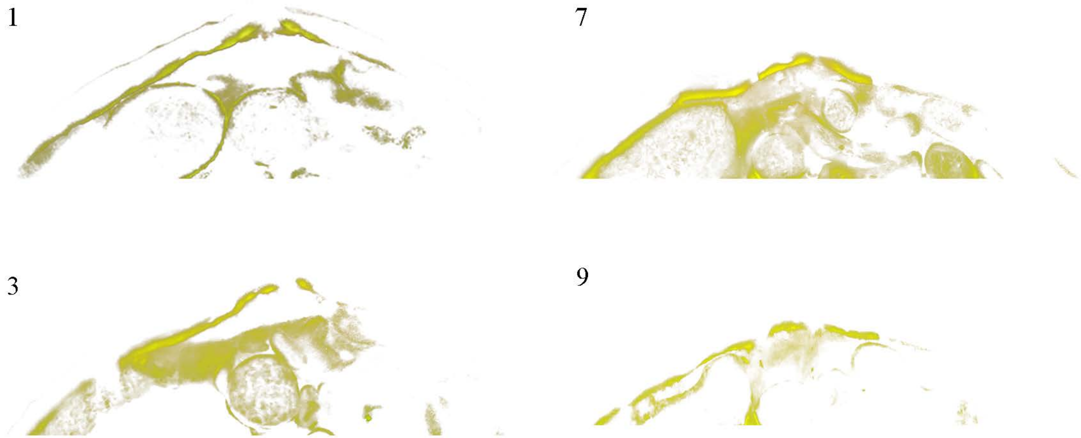


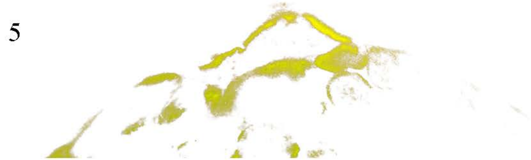


**Supplementary Fig. 1 Fluorescence signals of each fluorescence image extracted by Photoshop.** The images represent the slices 1, 3, 5, 7, and 9 (a). The fluorescence signals in the image was selected using color range by Photoshop cc, and the fuzziness for all images was uniformly set to 74. The selected fluorescence signals were save as a new layer, and the image was exported after changing the color to fcff00.

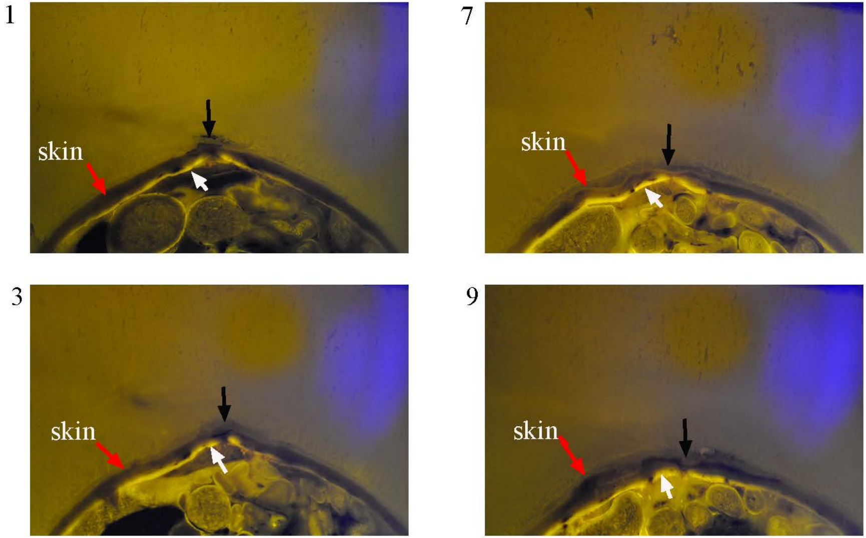


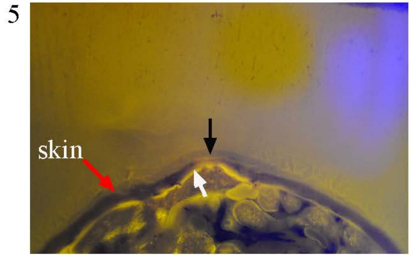


**Supplementary Fig. 2 Fluorescence image of each transverse sections along the ventral midline.** The images represent the slices 1, 3, 5, 7, and 9 (a). Black arrow indicates the position of ventral midline. White arrow indicates the fluorescence signals.

1

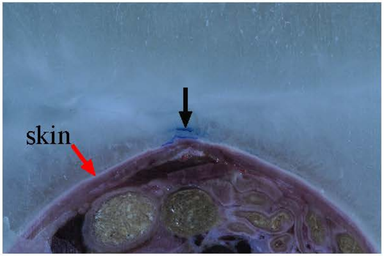


7

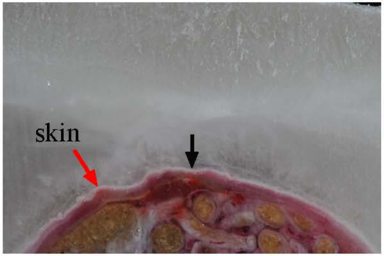


3

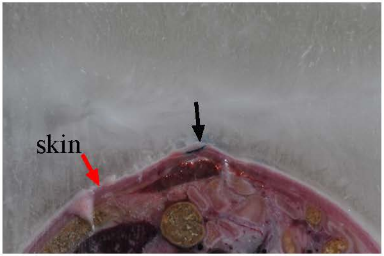


9

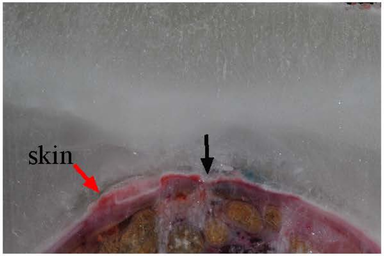


5

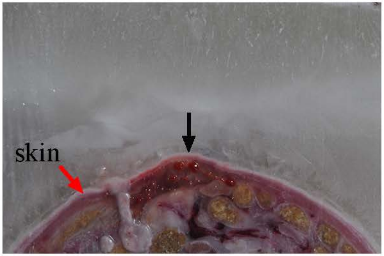


**Supplementary Fig. 3 Bright field image of each transverse sections along the ventral midline**.

The images represent the slices 1, 3, 5, 7, and 9 (a). Black arrow indicates the position of ventral midline.
